# Hunchback activates Bicoid in post-mitotic Pair1 neurons to regulate synapse number

**DOI:** 10.1101/2021.11.29.470460

**Authors:** Kristen M Lee, Amanda M Linskens, Chris Q Doe

## Abstract

The proper formation and function of neural circuits is crucial for cognition, sensation, and behavior. Neural circuits are highly-specific, and this specificity is dependent on neurons developing key features of their individual identities: morphology, anatomical location, molecular expression and biophysiological properties. Previous research has demonstrated that a neurons identity is, in part, generated by the temporal transcription window the neuron is born in, and the homeodomain transcription factors expressed in the mature neuron. However, whether temporal transcription factors and homeodomain transcription factors regulate neural circuit formation, maintenance and function remains unknown. Here, we utilize a well-characterized neural circuit in the *Drosophila* larvae, the Pair1 neuron. We determined that in the Pair1 neuron, the temporal transcription factor Hunchback activates the homeodomain transcription factor Bicoid (Bcd). Both Hunchback and Bcd are expressed in Pair1 throughout larval development. Interestingly, Hunchback and Bcd were not required in Pair1 for neurotransmitter identity or axonal morphology, but were required for synapse density. We found that these transcription factors were functioning post-mitotically in Pair1 to regulate synapse density. Additionally, knocking down Hunchback and Bcd in Pair1 neurons disrupted the behavioral output of the circuit. We utilized the genetic tool TransTango to determine that Hunchback function in Pair1 is to repress forming synapses with erroneous neurons. To our knowledge, these data are the first to show Hunchback activating Bcd expression, as well as the first to demonstrate a role for Hunchback and Bcd post-mitotically.

## Introduction

In the central nervous system, individual neurons connect and communicate with one another through synapses to form neural circuits^1^. Throughout life, the combined activity of neurons within circuits contributes to all nervous system function, including the generation of behavior. Aberrant neural circuit development has been associated with many CNS disorders, such as autism and ADHD^2^. Neuronal identity is defined by the anatomical, morphological, molecular and biophysiological properties of a neuron. Although neuronal molecular identity and morphology are well characterized, little is known about the mechanisms establishing neuronal synapse number, position and connectivity, key aspects of neuronal identity.

In Drosophila, embryonic neuronal identity is initially specified by the combination of spatial and temporal transcription factors (TFs). Spatial patterning genes create different neural stem cell (neuroblast) molecular identity^3^, followed by each neuroblast sequentially expressing a series of “temporal” TFs: Hunchback > Kruppel > Pdm (FlyBase: nubbin/pdm-2^4^) > Castor, which diversify neurons made from each neuroblast lineage^5^. Temporal TFs are known to specify axon and dendrite morphology and targeting^6–8^ as well as behavior^8^. For example, in neuroblast 7-1, the best characterized lineage in the embryo, the zinc-finger temporal TF Hunchback promotes expression of the homeodomain TF Even-skipped which is required for proper dorsal projecting motor neuron morphology and connectivity^9–12^; and the combination of Kruppel and Pdm temporal TFs promotes expression of the homeodomain TF Nkx6 (FlyBase: HGTX^4^) which is required for proper ventral projecting motor neuron morphology and connectivity^6,13^. In both cases, transient temporal TF expression activates a homeodomain TF which persists in the post-mitotic neuron to regulate neuron morphology. Similarly, work from the Hobert lab in C. elegans supports a model in which all neuronal identity is specified by combinations of homeodomain TFs^14^. Overall, from worms to flies to mammals, homeodomain TFs are known to specify molecular and morphological neuronal identity^15,16,17^.

Although homeodomain TFs are well known to specify these early aspects of neuronal identity^15,16,17^, their role in specifying later aspects of neuronal identity such as synapse number, position, and connectivity remains poorly understood. To address this question, we utilized the Pair1 locomotor circuit in *Drosophila*. The Pair1 neuron is a descending, GABAergic interneuron with ipsilateral dendrites and contralateral axonal projections^18–20^. The Moonwalker Descending Neurons (MDN) provide inputs to Pair1, and Pair1 sends outputs to A27h neurons in the ventral nerve cord (VNC)^18–20^. When optogenetically activated, the Pair1 neurons induce a pause in forward locomotion by inhibiting the A27h neurons, which drive forward locomotion^18–20^. Importantly, we previously reported that the temporal TF Hunchback and the homeodomain TF Bicoid (Bcd) are expressed in Pair1 throughout life^18^, providing candidates to study the transcriptional regulation of Pair1 neuronal identity and connectivity.

Hunchback is the first temporal TF to be expressed in the *Drosophila* embryo and gives rise to early born neurons^5^. In the ventral nerve cord (VNC), Hunchback is not required to maintain the identity of early born neurons post-mitotically^21^. Although Bcd is a homeodomain TF, its role outside of the early embryo remains unknown. In the early embryo, Bcd is a morphogen that directly activates transcription of *hb*^22^, which has a well-characterized role in specifying anterior structures along the anterior-posterior body axis^23^. Although Hunchback and its role in temporal patterning is conserved in mammals^24^, Bcd is found only in higher dipteran insects, making it an interesting contributor to insect evolution^25,26^.

Bcd directly activates *hunchback* transcription in the early embryo, raising the possibility that Bcd might have a similar function in the Pair1 neuron, where both Bcd and Hunchback are expressed^18^. Interestingly, we find the opposite genetic relationship between Bcd and Hunchback in the Pair1 neuron: Hunchback activates *bcd* expression. We go on to show that Hunchback or Bcd RNAi knockdown specifically in the Pair1 neuron results in ectopic synapse formation in Pair1, the formation of new patterns of connectivity, and a change in locomotor behavior. These are the first phenotypes for Hunchback and Bcd in post-mitotic neurons, and among the first temporal TF or homeodomain TF role in central synapse number and connectivity. Our data support the emerging model that temporal TFs drive expression of homeodomain TFs that go on to regulate multiple aspects of neuronal identity including synapse number/position, connectivity, and behavior.

## Results

### Hunchback promotes Bicoid expression in Pair1

Our previous research demonstrated that Pair1 neurons express Hunchback and Bcd during larval development and during adulthood^18^. In the embryonic blastoderm, Bcd activates Hunchback^23^. Here we ask whether Bcd activates Hunchback expression in Pair1 neurons. We labeled Pair1 neurons with GFP and visualized Hunchback and Bcd expression. Note that Bcd protein is detected in one or two cytoplasmic inclusions in the Pair1 neurons (seen with two independent antibodies and an FLAG-tagged Bcd protein; Figure 1A, Figure 3A-C, our previous work^18^, and data not shown). It is also seen in the nucleus at larval and adult stages^18^. Importantly, both nuclear and cytoplasmic Bcd are decreased by expression of Bcd RNAi (Figure 1B).

**Figure 1.**
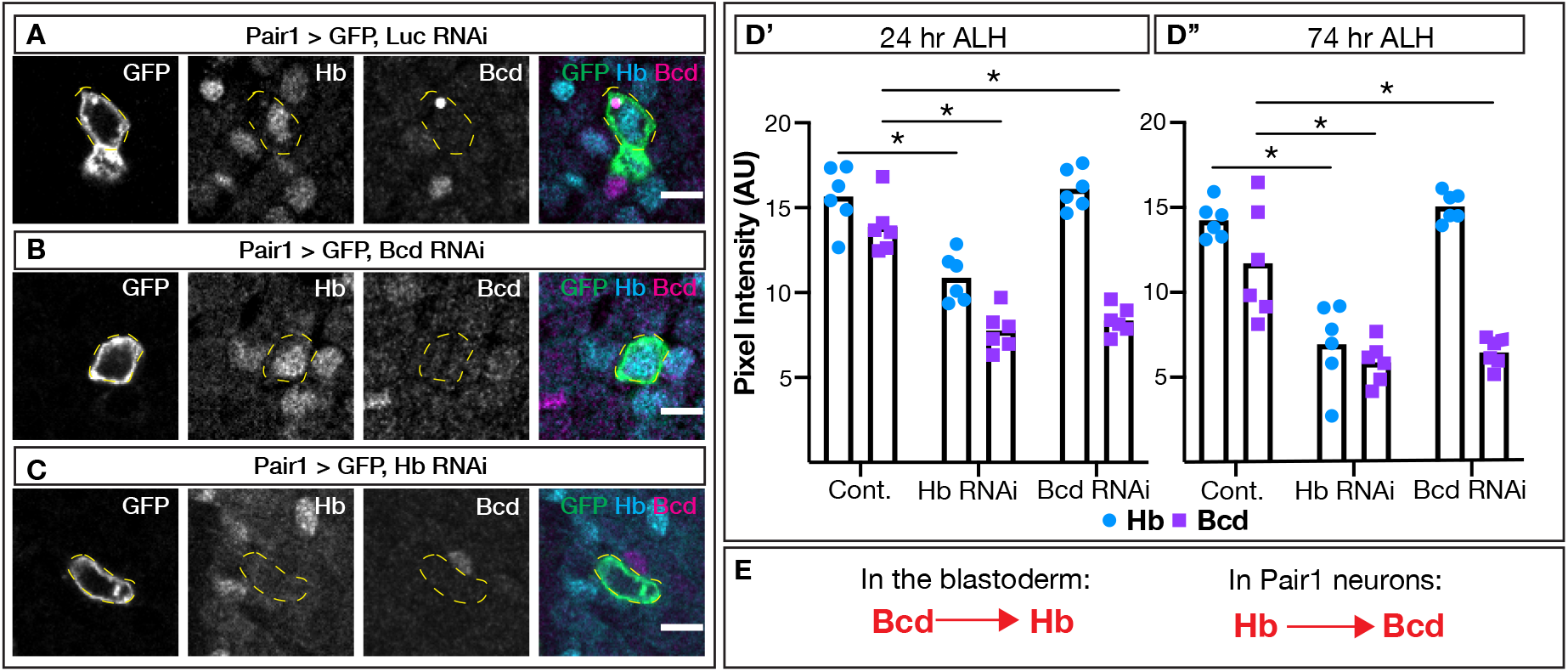
Hunchback activates Bicoid expression in larval Pair1 neurons. (A-C) Larval Pair1 neurons (GFP; far left column), Hunchback expression (Hb; middle left column), Bicoid expression (Bcd; middle right column) and merge (right column) at 24 hr after larval hatching (ALH). Scale bar, 5μm. (A) “Control”. Genotype: +*;UAS-myr::GFP; R75C02-Gal4/ UAS-Luc RNAi*, (B) “Bcd RNAi”. Genotype: +*;UAS-myr::GFP; R75C02-Gal4/ UAS-Bcd RNAi #2* (C) “Hb RNAi”. Genotype: +*;UAS-myr::GFP; R75C02-Gal4/ UAS-Hb RNAi*. (D) Quantification of Hb (blue) and Bcd (purple) expression within the Pair1 cell body at 24 hr ALH (D’) and 74 hr ALH (D”). Arbitrary units reported. (D’) Statistics: two-way ANOVA: genotype, F(2, 30) = 48.31, p < 0.001; protein, F(1, 30) = 86.69, p < 0.0001; interaction, F(2, 30) = 15.90, p < 0.0001; Bonferroni’s multiple comparisons between genotypes for each protein: Hb, cont vs Hb RNAi, p < 0.0001, cont vs Bcd RNAi, p > 0.9999; Bcd, cont vs Hb RNAi, p < 0.0001, cont vs Bcd RNAi, p < 0.0001; n = 6 animals. (D’’) Statistics: two-way ANOVA: genotype, F(2, 30) = 38.49, p < 0.001; protein, F(1, 30) = 42.78, p < 0.0001; interaction, F(2, 30) = 13.73, p < 0.0001; Bonferroni’s multiple comparisons between genotypes for each protein: Hb, cont vs Hb RNAi, p < 0.0001, cont vs Bcd RNAi, p = 0.9029; Bcd, cont vs Hb RNAi, p < 0.0001, cont vs Bcd RNAi, p < 0.0001; n = 6 animals. (E) Schematic of genetic interaction in early embryo (left) and larvae (right) between Hunchback and Bicoid.

In control animals expressing luciferase RNAi in Pair1, both Hunchback and Bcd show normal expression in Pair1 neurons (Figure 1A). In contrast, when the Bcd RNAi transgene was expressed in Pair1 neurons, Bcd expression was significantly reduced but Hunchback expression was unchanged (Figure 1B, 1D). Thus, Bcd does not activate Hunchback in Pair1. Next, we asked whether Hunchback could activate Bcd in Pair1 neurons. We expressed the Hunchback RNAi in Pair1 and observed both Hunchback and Bcd levels were significantly decreased (Figure 1C, 1D). We saw similar results at 24 and 74 hours after larval hatching (ALH; Figure 1D). We conclude that in Pair1 neurons, Hunchback activates Bcd expression, whereas Bcd does not activate Hunchback (Figure 1E). Thus, the genetic relationship between Hunchback and Bcd is the opposite in the early embryo and the Pair1 neuron.

### Hunchback is expressed in Pair1 throughout development

Hunchback is a early temporal TF expressed in the embryonic VNC and brain^5^. To determine if Pair1 neurons are born during the Hunchback temporal TF window, we labeled Pair1 with GFP and asked whether Pair1 neurons are expressed together with each of the embryonic temporal TFs in newly hatched larvae (0-4 hr ALH). Pair1 neurons expressed Hunchback and none of the other temporal TFs (Figure 2A). We previously showed that Hunchback was expressed in Pair1 neurons in the adult brain^18^, and here we extend these findings to show that Pair1 expresses Hunchback at all larval stages tested (6, 24, and 76 ALH; Figure 2B). We conclude that Pair1 is born during the Hunchback temporal window and is likely to maintain Hb expression into adulthood.

**Figure 2.**
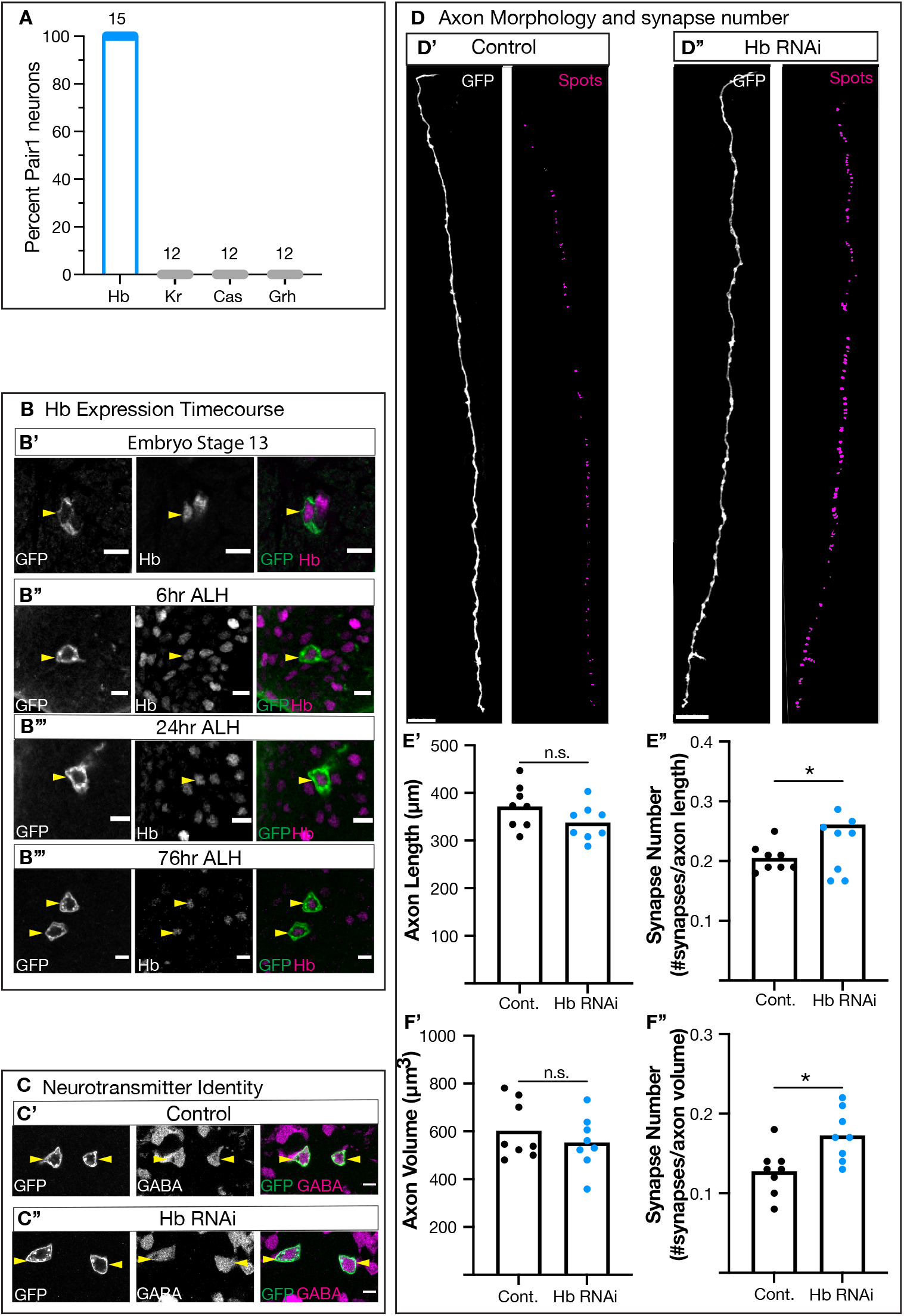
Hunchback expression in Pair1 regulates synapse number and position. (A) Percent of Pair1 neurons expressing Hunchback (Hb, left column, blue), Kruppel (Kr, grey, middle left column), castor (Cas, grey, middle right column) and grainy head (Grh, grey, right column). n = 12-15, reported for each protein. (B) Time course of Hunchback expression in Pair1 neurons. Pair1 neurons (GFP; left column), Hunchback (Hb; middle column) and merge (right column) at embryo stage 13 (B’), 6 hrs after larval hatching (ALH; B’’), 24 hr ALH (B’’’) and 76 hr ALH (B’’’’). Yellow arrows indicate Pair1 cell body. Scale bar, 5μm. Genotype: +*;UAS-myr::GFP; R75C02-Gal4* (C) Assessing GABA expression in Pair1 neurons. Pair1 neurons (GFP; left column), GABA expression (middle column) and merge (right column) in control animals (C’; Genotype: +*;UAS-myr::GFP; R75C02-Gal4/ UAS-Luc RNAi*) and Hunchback knockdown animals (C’’; Genotype: +*;UAS-myr::GFP; R75C02-Gal4/ UAS-Hb RNAi*). (D) Pair1 axons in control animals (D’) and hb knockdown animals (D’’). Pair1 axons (GFP, left column) and reconstructed axons (surface, green) and synapses (spots, magenta). Scale bar, 15 μm. (E’) Axon length in control (black) and *hb* RNAi (blue). Statistics: t-test, p = 0.1312, n = 8 animals. (E’’) Number of synapses normalized to axon length in control (black) and hb RNAi (blue). Statistics: t-test, p = 0.009, n = 8 animals. (F’) Axon volume in control (black) and hb RNAi (blue). Statistics: t-test, p = 0.4153, n = 8 animals. (F’’) Number of synapses normalized to axon volume in controls (black) and hb RNAi (blue). Statistics: t-test, p = 0.0117, n = 8 animals. (D-F) Control Genotype: *LexAop-myr::GFP; R75C02-LexA, LexAop-brp-Sh::mCherry/+; R75C02-Gal4/ UAS-Luc RNAi*. Hunchback RNAi Genotype: *LexAop-myr::GFP; R75C02-LexA, LexAop-brp-Sh::mCherry/+; R75C02-Gal4/ UAS-Hb RNAi*.

### Bicoid is expressed in Pair1 throughout development

We previously showed that Bicoid was expressed in Pair1 neurons in the adult brain^18^, but its expression at embryonic and larval stages has not been reported. We employed two complimentary strategies to assay for Bcd expression in Pair1 neurons. For both approaches, we labeled Pair1 neurons with GFP. In the first method, we imaged an epitope-tagged Bcd protein and found that it is expressed in Pair1 neurons at 6, 24 and 76 hrs ALH (Figure 3A). In the second method, we used two different antibodies against the Bcd protein and found that Bcd is expressed in the Pair1 cell body at 6, 24 and 76 hrs ALH (Figure 3B, 3C). For each Bcd antibody and the epitope-tagged protein, we observed the same cytoplasmic punctate sphere, which is absent following Bcd RNAi (see Figure 1). Our results, taken together with our previous results, show that Bcd is maintained in the Pair1 neurons from embryogenesis to adulthood, consistent with a model in which the. homeodomain Bcd TF acts to maintain some or all aspects of Pair1 neuronal identity.

**Figure 3.**
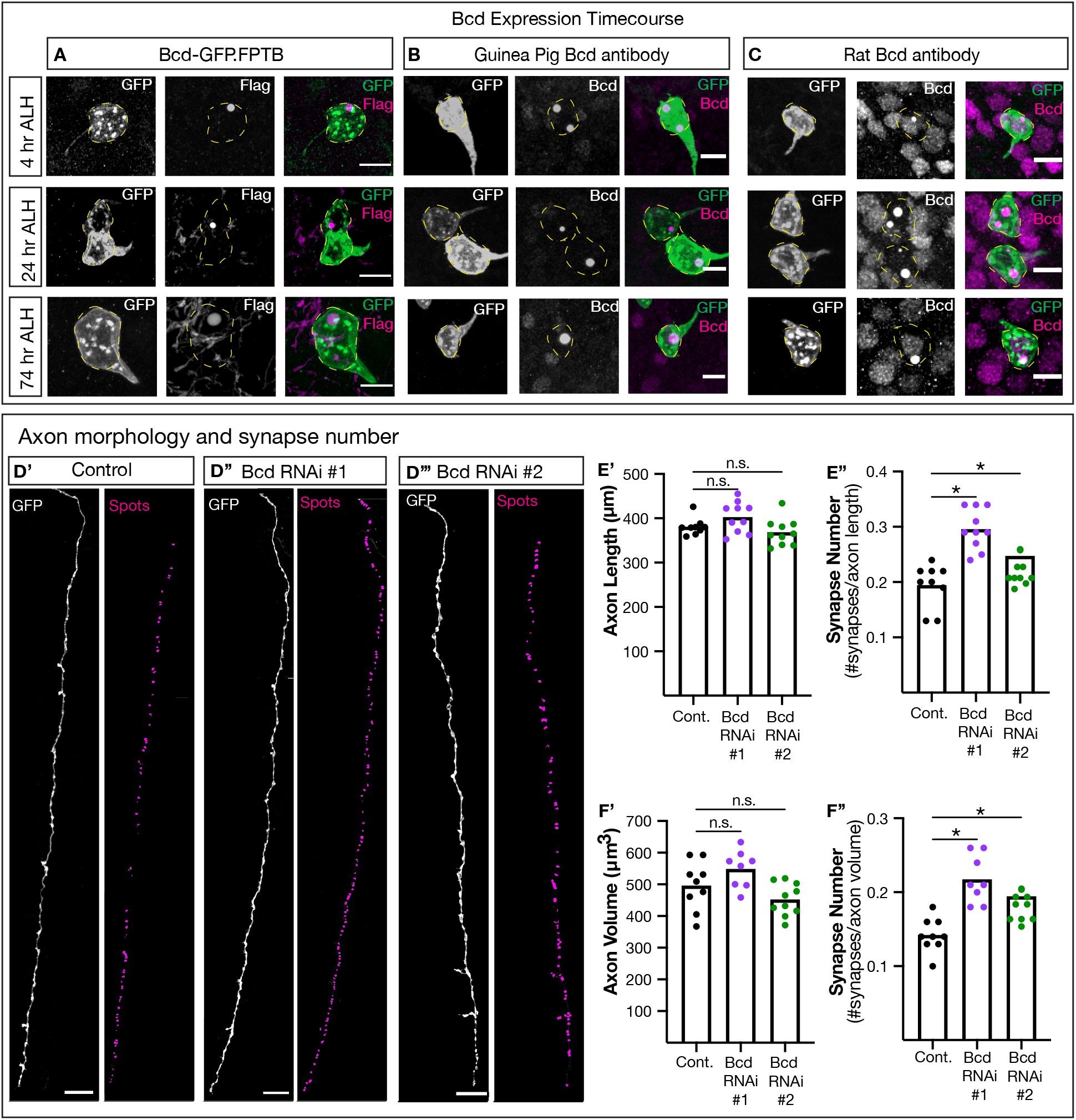
Bicoid expression in Pair1 regulates synapse number and position. (A-C) Independent bicoid labeling strategies in Pair1 neurons at 4 hrs ALH (top row), 24 hr ALH (middle row) and 74 hr ALH (bottom row). (A) Pair1 neurons (GFP, left column), Bcd tagged protein (Flag, middle column) and merge (right column). Scale bar, 5μm. Genotype: + *;UAS-myr::GFP/bcd-GFP.FPTB; R75C02-Gal4*. (B-C) Pair1 neurons (GFP, left column), Bicoid (Bcd, middle column) and merge (right column). Two independent Bcd primary antibodies used. Scale bar, 5μm. Genotype: + *;UAS-myr::GFP; R75C02-Gal4*. (D) Pair1 axons in control animals (D’) and Bicoid knockdown animals (D’’, D’’’). Pair1 axons (GFP, left column) and reconstructed axons (surface, green) and synapses (spots, magenta). Scale bar, 15 μm. (E’) Axon length in control (black), Bcd RNAi #1 (purple) and Bcd RNAi #2 (green) animals. Statistics: One-way ANOVA, F(2, 26) = 3.602, p = 0.0416; Bonferonni’s multiple comparisons, Cont vs Bcd RNAi #1, p = 0.2212, Cont vs Bcd RNAi #2, p = 0.7222; n = 9-10 animals. (E’’) Number of synapses normalized to axon length in controls (back), Bcd RNAi #1 (purple) and Bcd RNAi #2 (green) animals. Statistics: One-way ANOVA, F(2, 26) = 15.29, p < 0.0001; Bonferonni’s multiple comparisons, Cont vs Bcd RNAi #1, p < 0.0001, Cont vs Bcd RNAi #2, p = 0.0127; n = 9-10 animals. (F’) Axon volume in control (black), Bcd RNAi #1 (purple) and Bcd RNAi #2 (green) animals. Statistics: One-way ANOVA, F(2, 24) = 5.192, p = 0.0134; Bonferonni’s multiple comparisons, Cont vs Bcd RNAi #1, p = 0.1985, Cont vs Bcd RNAi #2, p = 0.2867; n = 8-10 animals. (F’’) Number of synapses normalized to axon volume in control (black), Bcd RNAi #1 (purple) and Bcd RNAi #2 (green) animals. Statistics: One-way ANOVA, F(2, 24) = 13.92, p < 0.0001; Bonferonni’s multiple comparisons, Cont vs Bcd RNAi #1, p < 0.0001, Cont vs Bcd RNAi #2, p = 0.0022; n = 8-10 animals. (D-F) Control Genotype: *LexAop-myr::GFP; R75C02-LexA, LexAop-brp-Sh::mCherry/+; R75C02-Gal4/ UAS-Luc RNAi*. Bcd RNAi #1 Genotype: *LexAop-myr::GFP; R75C02-LexA*, *LexAop-brp-Sh::mCherry*/+; *R75C02-Gal4/ UAS-Bcd RNAi #1*. Bcd RNAi #2 Genotype: *LexAop-myr::GFP; R75C02-LexA, LexAop-brp-Sh::mCherry/+; R75C02-Gal4/ UAS-Bcd RNAi #2*.

### Hunchback and Bcd are not required for Pair1 neurotransmitter identity or axon morphology

Here we test the role of Hunchback in establishing or maintaining neurotransmitter identity and axon morphology in the Pair1 neuron. Pair1 expresses the neurotransmitter GABA and sends contralateral axons down the entirety of the larval VNC. We knocked down Hunchback specifically in Pair1 using the validated Hunchback RNAi transgene (Figure 1) and screened for GABA expression and axon morphology at 76 hours ALH. The Pair1-Gal4 driver does not label any additional neurons at 76 hrs, making it an ideal timepoint for this analysis. We found that loss of Hunchback in the Pair1 neuron had no detectable effect on GABA levels in Pair1 (Figure 2C). Next, we assessed axon morphology by measuring axon length and volume using Imaris image analysis software (Figure 2D, “Surface”). Compared to control, when Hunchback was knocked down axon length and volume was not changed (Figure 2E, 2F). Because loss of Hunchback eliminates Bcd in Pair1 (see above), we conclude that reduction in both Hunchback and Bcd has no effect on Pair1 neurotransmitter expression or axon morphology.

Although Hunchback knockdown reduces Bcd level, and thus it is likely that neither Hunchback or Bcd are required for axon morphology, we wanted to confirm this by directly knocking down Bcd. Therefore, we knocked down Bcd using two independent RNAi transgenes in Pair1 neurons and assayed axon morphology, as described above. Compared to control, when Bcd was knocked down axon length and volume was not changed (Figure 3D-F). We conclude that neither Hunchback nor Bcd is required for Pair1 axon morphology.

### Hunchback and Bcd are required for Pair1 synapse number and position

Next we assayed Hunchback RNAi knockdown for changes in synapse number and position. We expressed the pre-synaptic non-functional tag Bruchpilot-Short (Brp) in Pair1 and quantified Brp+ puncta along the Pair1 axons using Imaris (Figure 2D, “Spots”) to get synapse number. To calculate synapse density, we normalized the number of synapses to the axon length and volume. We found that compared to controls, Hunchback knockdown led to a significant increase in synapse number (Figure 2E, 2F). Similarly, Bcd knockdown significantly increased synapse number compared to control (Figure 3D-F). Interestingly, the ectopic synapses were preferentially localized to the thoracic region of the Pair1 neuron (data not shown). The similarity of the Bcd and Hunchback phenotypes suggest that Hunchback acts indirectly, by activating Bcd, to maintain synapse numbers, whereas Bcd is likely to act more directly to regulate synapse numbers (see Discussion). We conclude that Hunchback and Bcd are required to stabilize Pair1 synapse number by preventing the formation of pre-synapses in the thoracic region of the Pair1 neuron. Note that both Hunchback and Bcd knockdown are performed constitutively beginning in the embryo, so we can’t make any conclusion about when during development that Hunchback and Bcd act to regulate synapse number.

### Hunchback and Bicoid act in post-mitotic Pair1 neurons to regulate synapse number

To date, Hunchback and Bcd function has been primarily investigated in the developing embryo^5,23^. To investigate a novel role for Hunchback and Bcd post-mitotically, we utilized Gal80ts to knockdown Hunchback and Bcd at embryo Stage 16, when Pair1 neurons are post-mitotic. To accomplish this, we reared animals with the Pair1-Gal4, Gal80ts and Hunchback RNAi transgenes at 18C; at this temperature the Gal80ts protein is in an active conformation allowing the Gal80 to inhibit Gal4 activation of the RNAi transgene. At embryonic stage 16, the animals were switched to 30; at this temperature the Gal80ts protein is in an inactive conformation allowing the Gal4 to drive expression of the RNAi transgene^27^ (Figure 4A). At 76hr ALH we assayed Pair1 axonal volume, length, and synapse density as described previously. When Hunchback and Bcd were individually knocked down in post-mitotic Pair1 neurons, axon volume and length was not altered compared to control (Figure 4B-F), similar to our constitutive Hunchback and Bcd knockdown (see Figure 2, 3). However, Pair1 neuron synapse number was significantly increased when Hunchback and Bcd were individually knocked down in post-mitotic Pair1 neurons (Figure 4B-F). This indicates that Hunchback and Bcd can function post-mitotically. This result is also the first example of the temporal TF Hunchback having a function in post-mitotic neurons, and the first example that the homeodomain TF Bcd has a role beyond the early embryo.

**Figure 4.**
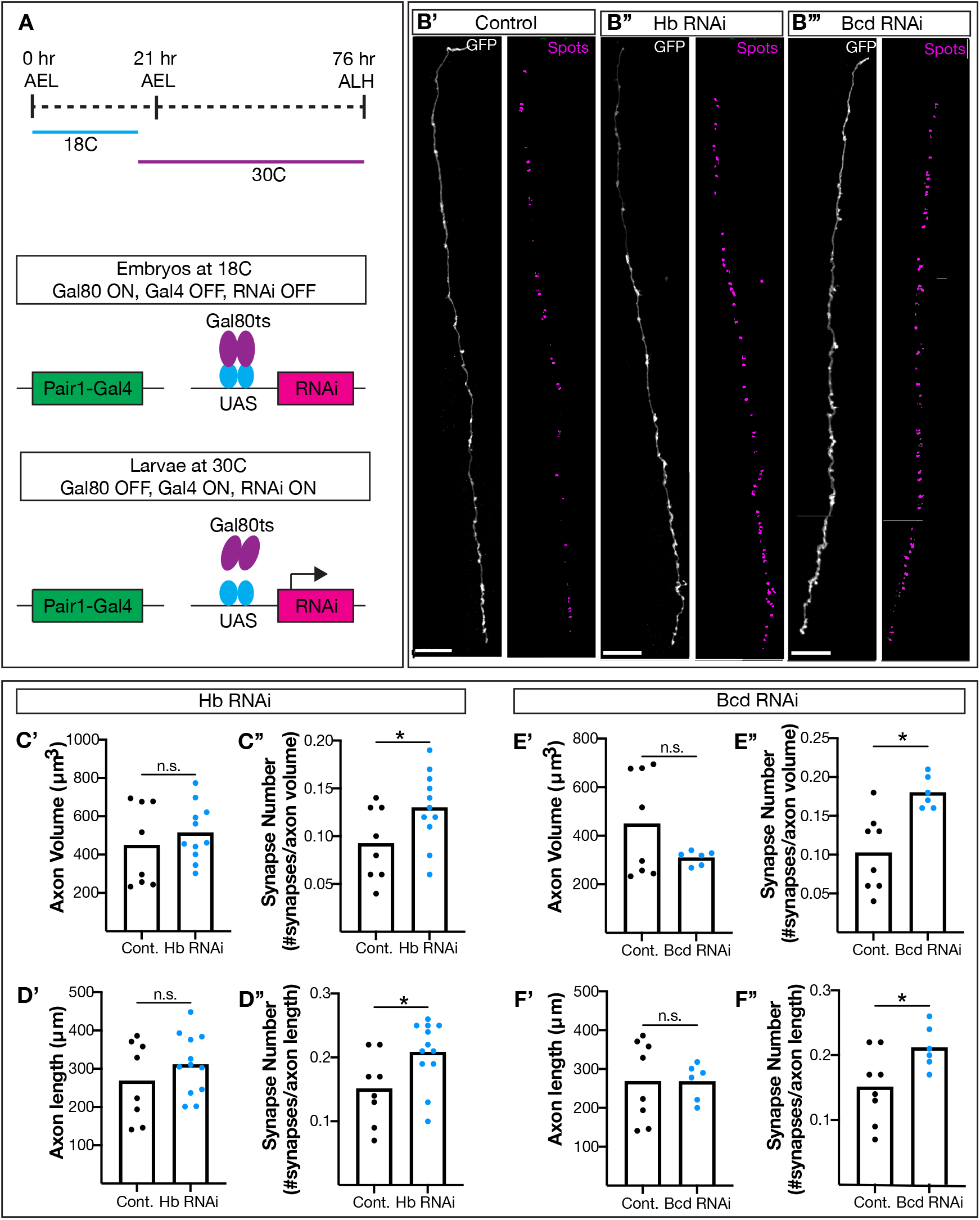
Hunchback and Bicoid function post-mitotically in Pair1 neurons. (A) Schematic of the experimental design. (B) Pair1 axons in control animals (B’), Hb RNAi (B’’) and Bcd RNAi (B”’) animals. Pair1 axons (GFP, left column) and reconstructed axons (surface, green) and synapses (spots, magenta). Scale bar, 20 μm. (C-D) Hb knockdown in Pair1 neurons. Control Genotype: *LexAop-myr::GFP; R75C02-LexA, LexAop-brp-Sh::mCherry/Gal80ts; R75C02-Gal4/ UAS-Luc RNAi*. Hb RNAi Genotype: *LexAop-myr::GFP; R75C02-LexA, LexAop-brp-Sh::mCherry/Gal80ts; R75C02-Gal4/ UAS-Hb RNAi*. (C’) Axon volume in control (black) and Hb RNAi (blue) animals. Statistics: t-test, p = 0.4432, n = 8-11. (C’’) Number of synapses normalized to axon volume in control (black) and Hb RNAi (blue) animals. Statistics: t-test, p = 0.0495, n = 8-11. (D’) Axon length in control (black) and Hb RNAi (blue) animals. Statistics: t-test, p = 0.3115, n = 8-12. (D’’) Number of synapses normalized to axon length in control (black) and Hb RNAi (blue) animals. Statistics: t-test, p = 0.0271, n = 8-12. (E-F) Bcd knockdown in Pair1 neurons. Bcd RNAi Genotype: Control Genotype: *LexAop-myr::GFP; R75C02-LexA, LexAop-brp-Sh::mCherry/Gal80ts; R75C02-Gal4/ UAS-Bcd RNAi #2*. (E’) Axon volume in control (black) and Bcd RNAi (blue) animals. Statistics: t-test, p = 0.1393, n = 6-8. (E’’) Number of synapses normalized to axon volume in control (black) and Bcd RNAi (blue) animals. Statistics: t-test, p = 0.0037, n = 6-8. (F’) Axon length in control (black) and Bcd RNAi (blue) animals. Statistics: t-test, p = 0.9914, n = 6-8. (F’’) Number of synapses normalized to axon length in control (black) and Bcd RNAi (blue) animals. Statistics: t-test, p = 0.0349, n = 6-8.

#### Hunchback and Bicoid are required in Pair1 for normal locomotion

It is well-characterized that when the Pair1 neuron is optogenetically activated the larvae pause^18–20^. Since knocking down Hunchback and Bcd in Pair1 neurons results in extra synapses, we wanted to know whether the extra synapses disrupt overall circuit function. To investigate this, we knocked down Hunchback and Bcd individually in Pair1 neurons and assayed locomotion before, during, and after activation of Pair1 by the red light gated cation channel CsChrimson^19^. First, we knocked down Hunchback in Pair1 and assayed larval speed (Figure 5A). We observed no differences in the speeds of control or Hunchback knockdown larvae before the red light stimulus was presented. However, upon Pair1 activation, the Hunchback knockdown larvae showed faster pausing, measured by a significantly steeper negative slope (Figure 5A’). We observed that control animals quickly increase their speeds after the initial pause during red light exposure, whereas the Hunchback knockdown animals remained slow. To quantify this, we normalized the speeds of the control and Hunchback knockdown animals during red light exposure to the average control speed. We found that Hunchback knockdown in Pair1 resulted in a significantly slower speed while the red light stimulus was presented (Figure 5A”). Lastly, we observed that after the red light exposure, control animals quickly return to a baseline speed (speed before red light stimuli), while the Hunchback knockdown animals took longer. To quantify this, we normalized the speeds of the control and Hunchback knockdown animals after red light exposure to the average control speed. We found that Hunchback knockdown in Pair1 resulted in a significantly slower speed after red light exposure compared to control (Figure 5A”’). We conclude that Hunchback is required in Pair1 for normal Pair1-dependent locomotion.

**Figure 5.**
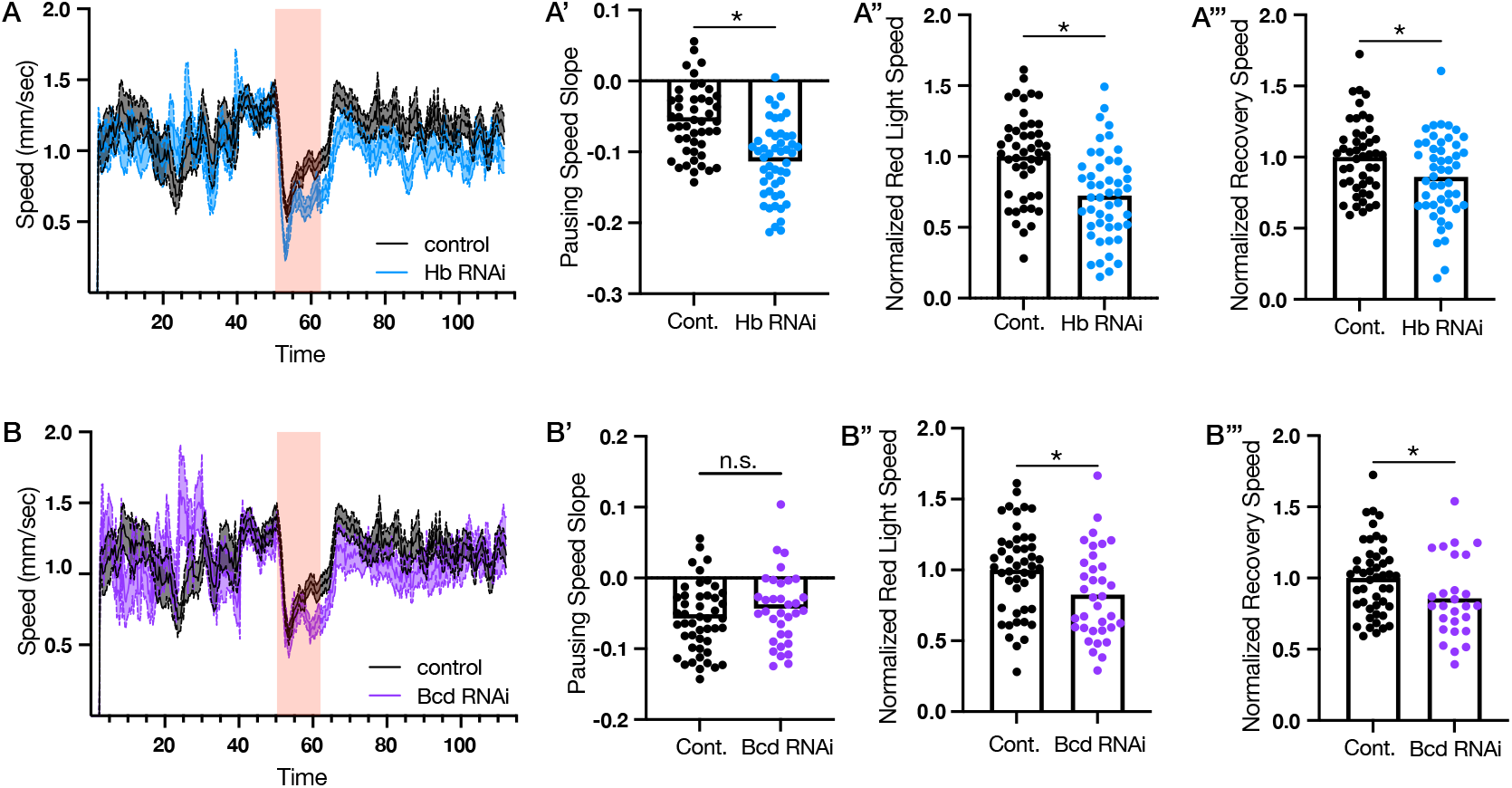
Hunchback and Bicoid regulate Pair1-dependent locomotor behavior. (A) Speed of control (black) and Hb RNAi (blue) larvae over time. Red bar represents red light exposure. (A’) Pausing speed slope in control (blue) and Hb RNAi (blue) animals. Statistics: t-test, p < 0.0001, n = 47-49 animals. (A’’) Normalized speed during red light exposure of control (black) and Hb RNAi (blue) animals. Statistics: t-test, p < 0.0001, n = 47-49 animals. (A’’’) Normalized speed during after the red light stimulus in control (black) and Hb RNAi (blue) animals. Statistics: t-test, p = 0.0153, n = 47-49. Control Genotype: *UAS-CsChrimson::mVenus;;R75C02-Gal4/UAS-Luc RNAi*. Hb RNAi Genotype: *UAS-CsChrimson::mVenus;;R75C02-Gal4/UAS-Hb RNAi*. (B) Speed of control (black) and Hb RNAi (purple) larvae over time. Red bar represents red light exposure. (B’) Pausing speed slope in control (blue) and Bcd RNAi (purple) animals. Statistics: t-test, p = 0.2091, n = 34-47 animals. (B’’) Normalized speed during red light exposure of control (black) and Bcd RNAi (purple) animals. Statistics: t-test, p = 0.0145, n = 34-47 animals. (B’’’) Normalized speed during after the red light stimulus in control (black) and Bcd RNAi (purple) animals. Statistics: t-test, p = 0.0304, n = 34-47. Bcd RNAi Genotype: *UAS-CsChrimson::mVenus;;R75C02-Gal4/UAS-Bcd RNAi #2*.

To determine if the previous phenotype was due to loss of Bcd in the Hunchback RNAi knockdown larvae, we knocked down Bcd in Pair1, assayed locomotion and performed the same quantifications. When Bcd was knocked down in Pair1, there was no change in pausing speed during Pair1 activation, but there was a significant decrease in overall speed during red light stimulus and the overall speed after the red light exposure (Figure 5B). Taken together, these data suggest that Hunchback and Bcd are required in Pair1 neurons for overall Pair1 neuronal circuit function.

### Hunchback regulates Pair1 connectivity

When activated, the Pair1 neurons inhibit their downstream partners, A27h, which normally induce forward locomotion^19^. Since Hunchback and Bcd are required in Pair1 neurons to maintain synapse number and regulate circuit function, we wanted to determine whether the disruptions in circuit function were due to either (i) A27h neurons forming more connections with Pair1 or (ii) new neurons forming connections with Pair1. Since either of these hypotheses could explain the behavioral phenotypes observed, we utilized TransTango which is a technique to label partners downstream of Pair1 with an HA epitope tag^28^. We focused our experiments on Hunchback. We expressed the TransTango transgene and Hunchback RNAi transgene in Pair1 neurons, counted the number of HA+ cell bodies labeled downstream of Pair1, and compared these results to controls with normal Hunchback levels. To unbiasedly differentiate the VNC (where Pair1-A27h synapses are located), from the central brain lobes, we labeled the suboesophageal zone with the Sex Combs Reduced (Scr) antibody^29^ and only counted cell bodies posterior of the Scr staining (dashed line, Figure 6). Comparing our control Pair1>TransTango data with the first instar larval TEM volume, both show that a majority of Pair1 synaptic partners are in the posterior region of the VNC^19,20^, validating the TransTango results. We found that knocking down Hunchback in Pair1 neurons significantly increased the number of neurons labeled by the TransTango transgene (Figure 6). This shows that new neurons are connecting to Pair1, although we can’t rule out the possibility that there are also more Pair1-A27h synapses. We conclude that Hunchback (and likely Bcd) are required for proper Pair1 connectivity.

**Figure 6.**
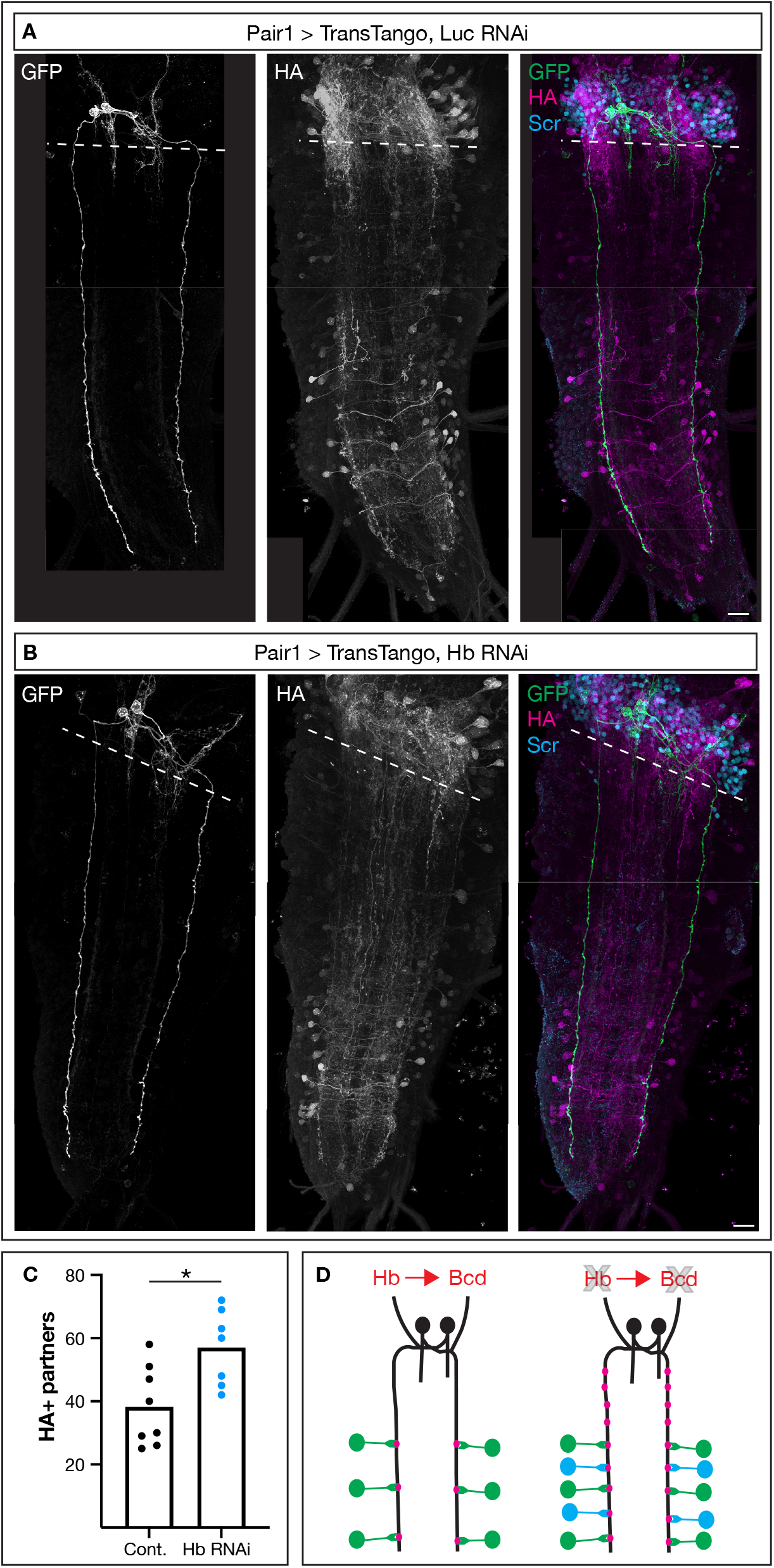
Hunchback regulates Pair1 connectivity. (A, B) Utilizing the TransTango genetic tool to visualize Pair1 neurons (GFP, left column), neuronal partners downstream of Pair1 (HA, middle column) and merge (right column). Dash line indicates the posterior edge of the Scr+ (cyan) region. Scale bar, 15 μm. (A) Control animals. Genotype: *UAS-myr::GFP.QUAS-mtdTomato-3XHA; P(trans-Tango); R75C02-Gal4/UAS-Luc RNAi*. (B) Hb knockdown. Genotype: *UAS-myr::GFP.QUAS-mtdTomato-3XHA; P(trans-Tango); R75C02-Gal4/UAS-Hb RNAi*. (C) Quantification of HA+ cell bodies posterior to the Scr+ region in control (black) and Hb RNAi (blue) animals. Statistics: t-test, p = 0.0015, n = 7-8. (D) Schematic demonstrating Hunchbacks role in mediating both synapse number and synaptic partner selection in Pair1 neurons.

## Discussion

Our results show that in larvae, Hunchback activates Bcd in the post-mitotic Pair1 neuron, where it is required for maintaining synapse number, synapse density, and connectivity. When Hunchback or Bcd levels are decreased, synapse density is increased, with a corresponding disruption of the function of the Pair1 locomotor neural circuit. This disruption is likely through the Pair1 neuron synapsing with new neuronal partners. This work demonstrates a novel role for Hunchback and Bcd – functioning post-mitotically to maintain neural circuits.

Unlike most early-born neurons in the VNC that only transiently express Hunchback^21^, and Bcd which is only expressed in the first few hours of embryogenesis, the Pair1 neuron maintains both Hunchback and Bcd expression into the adult. This suggests that a Pair1-specific regulatory mechanism may be leading to the persistent Hunchback and Bcd expression and function. Given that the Pair1 neuron persists into adulthood, still expresses Hunchback and functions within a similar locomotor neural circuit^18^, we hypothesize that Hunchback and Bcd expression may be required in Pair1 neurons throughout life for the maintenance of the Pair1 locomotor neural circuit.

Previously, it has been shown that Bcd activates *hunchback* in the early embryo^23^. Our study is the first we are aware of to demonstrate the reverse: that Hunchback can activate *bcd in vivo*. This finding supports our initial hypothesis that temporal transcription factors, like Hunchback, can activate homeodomain transcription factors, like Bcd, to specify some or all aspects of neuronal identity. We found that Hunchback and Bcd had no detectable role in regulating axon morphology or neurotransmitter expression, key aspects of neuronal identity. Yet we found both Hunchback and Bcd to be required for maintaining synapse number and appropriate connectivity in the Pair1 neuron. Thus, Hunchback and Bcd are required for some but not all aspects of neuronal identity.

Interestingly, it appears that Bcd is not the only homeodomain TF functioning downstream of Hunchback in Pair1. When Hunchback was knocked down in Pair1, pausing speed was increased and recovery speeds were decreased. However, Bcd knockdown only replicated the decreased recovery speed phenotype (Figure 5); this suggests that another homeodomain TF may be functioning downstream of Hunchback to regulate pausing speed. It is interesting to speculate that this transcription factor may be mediating synapses specifically with A27h, leading to more A27h inhibition when Pair1 is optogenetically activated and therefore a more rapid pause. On the other hand, Bcd could be functioning to inhibit connectivity with other neurons. Therefore, loss of Bcd would lead to novel neuronal partners, thus disrupting the recovery of the locomotor circuit after Pair1 activation. The data presented here begin to support this hypothesis, but additional work is needed to identify other homeodomain TFs functioning downstream of Hunchback.

To our surprise, primary Bcd protein expression in Pair1 neurons was a spherical puncta located in the cytoplasm; this was observed with two independent Bcd antibodies and a third FLAG-tagged Bcd protein, and abolished by Bcd RNAi. The spherical puncta may represent a phase separation condensate^30^. In addition, we also note that each labeling method showed different levels of nuclear staining in neurons surrounding Pair1^18^. Further investigation is needed to understand nature of the Bcd cytoplasmic puncta, but these studies have the potential to elucidate a novel role for phase separation in neurons.

Our work is the first, to our knowledge, to demonstrate a role for Hunchback and Bcd in post-mitotic neurons to regulate synapse number, connectivity, and circuit function. Our results raise the question of which is the more ancestral function of these two TFs: in segmentation, temporal patterning in neuroblasts, or post-mitotic neuron circuit formation?

## Materials and Methods

### Fly Husbandry

All flies were reared in a 25C room at 50% relative humidity with a 12 hr light/dark cycle unless noted otherwise. All comparisons between groups were based on studies with flies grown, handled, and tested together.

### Fly Stocks

Stocks:

- R75C02-Gal4 (referred to as Pair1-Gal4), BDSC #39886
- R75C02-LexA, BDSC #54365
- ;UAS-myr::GFP, BDSC #32198
- LexAop-myr::GFP, BDSC #32211
- UAS-Luc RNAi, BDSC #31603
- UAS-Hunchback RNAi, BDSC #34704
- UAS-Bcd RNAi #1 attp2, BDSC #33886
- UAS-Bcd RNAi #2 attp2, BDSC #35478
- Bcd-GFP.FPTB, BDSC #67654
- LexAop-brp-Sh::mCherry
- Gal80ts, BDSC #7019
- UAS-CsChrimson::mVenus, Gift from Vivek Jayaraman, JRC
- TransTango, BDSC #77124

### Immunostaining and Imaging

Standard confocal microscopy and immunocytochemistry methods were performed as previously described^19^. Primary antibodies used: Chicken anti-GFP (1:1500; Abcam, Eugene, OR) Rabbit anti-Hunchback (1:400; Doe Lab), Rabbit anti-GABA (1:500; Sigma, St. Louis, MO), Rabbit anti-mCherry (1:500; Novus, Centennial, CO), Goat anti-Bcd (1:1500; Gift from John Reinitz, University of Chicago, IL), Rat anti-Bcd (1:100; Gift from John Reinitz, University of Chicago, IL), Rat anti-Flag (1:400; Novus, Centennial, CO), Rat anti-HA (1:100; Sigma, St. Louis, MO) and Mouse anti-Scr (1:10; DSHB, Iowa City, IA). Secondary anitbodies were from Jackson ImmunoResearch (Donkey, 1:400; West Grove, PA). Confocal image stacks were acquired on a Zeiss 800 microscope. All images were process either in Fiji (https://imagej.new/fiji) or Imaris (https://imaris.oxinst). All figures were generated on Adobe Illustrator (Adobe, San Jose, CA).

### Quantification of Pixel Intensity

All pixel intensity quantification was done manually using the “Measure” feature in Fiji. The freehand tool was used to outline the cell body. Measurements were set to “Area” and “Raw Integrated Density”. Each slice imaging the cell body was measured. For each slice, the “Raw Integrated Density” was divided by the “Area”. The sum of these values is reported in arbitrary units.

### Quantification of Axon Phenotypes

To quantify axon length, the filament tool was used in Imaris Image Analysis Software. A region of interest was placed over one Pair1 axon labeled via GFP. The largest diameter was set to 1.7 and the smallest diameter was set to 0.2.

To quantify axon volume, the surface tool was used in Imaris Image Analysis Software. A region of interest was placed over one Pair1 axon labeled via GFP. The surface detail was set to 0.18 and the background subtraction was set to 1.8. The final threshold was set to 7.7% of the total.

To quantify synapse number we used the spots tool was used in Imaris Image Analysis Software on the Bruchpilot-Short::mCherry label. The XY diameter was set to 0.7 and the Z diameter was set to 1.5. The final threshold was set to 0.5% of the total. Each spot associated with each surface was calculated, with the threshold set to 0. To calculate synapse density, the total number of Spot was divided by the Length or Volume of the axon. As axon volume was unchanged by all manipulations, synapse density also represents synapse number.

### Behavior

At 48 hr ALH, larvae were transferred to apple caps with 0.5 μM All Trans Retinol (ATR) or 0.5 μM ethanol (vehicle). All behavior was done with third instar larvae (72-76 hr ALH). At approximately 72 hr ALH, larvae were transferred to an 1.2% agar sheet on the FIM table^31^. After two minutes to acclimate, larvae locomotion was recorded at 4 frames per second for 112 seconds with a Basler camera. At 50 seconds, the red light was turned on for 12 seconds. Larvae were tracked using FimTrak^31^ and acceleration was reported. To report pausing speed slope, the slope ((y1-y2)/(x1-x2)) of the speed before red light stimuli to the minimal speed during the red light stimuli was calculated. y1 was the animals speed before the light stimulus, y2 was the animals minimal speed during the light stimulus, x1 was the time which y1 was calculated, x2 was the time at which the animal reached minimal speed during the red light stimulus. To calculate the normalized red light speed, the average speed during the red light stimuli was calculated for each animal for each genotype. For the control genotype, the average red light stimuli speed for all animals was determined, termed the total control average. Then, the average red light speed for each animal for each genotype was divided by the total control average. The same approach was used to calculate normalized recovery speed.

#### TransTango

We utilized the TransTango transgene as described previously^28^. For the analysis, all cell bodies posterior of the Scr stain was counted, using the “Cell Counter” plugin in Fiji.

### Statistics

All statistical analysis were performed with Prism 9 (GraphPad Software, San Diego, CA). Numerical data in graphs show individual measurements (dots) and means (bars). The number of replicates (n) and definition of measurement reporter (i.e. animal, axon) for each data set is in the corresponding legend.

## Acknowledgements

We thank John Reinitz for antibodies and Vivek Jayaraman for fly stocks. Stocks obtained from the Bloomington Drosophila Stock Center (NIH P40D018537) were used in this study. Funding was provided by HHMI (CQD).

## Author Contributions

KML conceptualized, curated data, analyzed data, validated results, wrote and edited the manuscript, and generated figures. AML curated data, analyzed data, generated figures and edited the manuscript. CQD conceptualized, supervised, and edited the manuscript and figures.

## References

1. Shirasaki, R. & Pfaff, S. L. Transcriptional codes and the control of neuronal identity. Annu Rev Neurosci 25, 251–281 (2002).

2. Hoxha, E. et al. Maturation, Refinement, and Serotonergic Modulation of Cerebellar Cortical Circuits in Normal Development and in Murine Models of Autism. Neural Plast 2017, 6595740 (2017).

3. Crews, S. T. Drosophila Embryonic CNS Development: Neurogenesis, Gliogenesis, Cell Fate, and Differentiation. Genetics 213, 1111–1144 (2019).

4. Larkin, A. et al. FlyBase: updates to the Drosophila melanogaster knowledge base. Nucleic Acids Research 49, D899–D907 (2021).

5. Doe, C. Q. Temporal Patterning in the Drosophila CNS. Annu Rev Cell Dev Biol 33, 219–240 (2017).

6. Seroka, A., Yazejian, R. M., Lai, S.-L. & Doe, C. Q. A novel temporal identity window generates alternating Eve+/Nkx6+ motor neuron subtypes in a single progenitor lineage. Neural Dev 15, 9 (2020).

7. Seroka, A. Q. & Doe, C. Q. The Hunchback temporal transcription factor determines motor neuron axon and dendrite targeting in Drosophila. Development 146, dev175570 (2019).

8. Sullivan, L. F., Warren, T. L. & Doe, C. Q. Temporal identity establishes columnar neuron morphology, connectivity, and function in a Drosophila navigation circuit. Elife 8, e43482 (2019).

9. Doe, C. Q., Smouse, D. & Goodman, C. S. Control of neuronal fate by the Drosophila segmentation gene even-skipped. Nature 333, 376–378 (1988).

10. Isshiki, T., Pearson, B., Holbrook, S. & Doe, C. Q. Drosophila neuroblasts sequentially express transcription factors which specify the temporal identity of their neuronal progeny. Cell 106, 511–521 (2001).

11. Landgraf, M., Jeffrey, V., Fujioka, M., Jaynes, J. B. & Bate, M. Embryonic origins of a motor system: motor dendrites form a myotopic map in Drosophila. PLoS Biol 1, E41 (2003).

12. Fujioka, M. et al. Even-skipped, acting as a repressor, regulates axonal projections in Drosophila. Development 130, 5385–5400 (2003).

13. Broihier, H. T., Moore, L. A., Van Doren, M., Newman, S. & Lehmann, R. zfh-1 is required for germ cell migration and gonadal mesoderm development in Drosophila. Development 125, 655–666 (1998).

14. Reilly, M. B., Cros, C., Varol, E., Yemini, E. & Hobert, O. Unique homeobox codes delineate all the neuron classes of C. elegans. Nature 584, 595–601 (2020).

15. Tursun, B., Patel, T., Kratsios, P. & Hobert, O. Direct conversion of C. elegans germ cells into specific neuron types. Science 331, 304–308 (2011).

16. Serrano-Saiz, E. et al. Modular control of glutamatergic neuronal identity in C. elegans by distinct homeodomain proteins. Cell 155, 659–673 (2013).

17. Narita, Y. & Rijli, F. M. Hox genes in neural patterning and circuit formation in the mouse hindbrain. Curr Top Dev Biol 88, 139–167 (2009).

18. Lee, K. & Doe, C. Q. A locomotor neural circuit persists and functions similarly in larvae and adult Drosophila. Elife 10, e69767 (2021).

19. Carreira-Rosario, A. et al. MDN brain descending neurons coordinately activate backward and inhibit forward locomotion. Elife 7, (2018).

20. Tastekin, I. et al. Sensorimotor pathway controlling stopping behavior during chemotaxis in the Drosophila melanogaster larva. Elife 7, (2018).

21. Hirono, K., Kohwi, M., Clark, M. Q., Heckscher, E. S. & Doe, C. Q. The Hunchback temporal transcription factor establishes, but is not required to maintain, early-born neuronal identity. Neural Dev 12, 1 (2017).

22. Porcher, A. & Dostatni, N. The bicoid morphogen system. Curr Biol 20, R249–254 (2010).

23. Simpson-Brose, M., Treisman, J. & Desplan, C. Synergy between the hb and bicoid morphogens is required for anterior patterning in Drosophila. Cell 78, 855–865 (1994).

24. Wernet, M. F., Huberman, A. D. & Desplan, C. So many pieces, one puzzle: cell type specification and visual circuitry in flies and mice. Genes Dev 28, 2565–2584 (2014).

25. Lynch, J. & Desplan, C. Evolution of development: beyond bicoid. Curr Biol 13, R557–559 (2003).

26. McGregor, A. P. How to get ahead: the origin, evolution and function of bicoid. Bioessays 27, 904–913 (2005).

27. McGuire, S. E., Roman, G. & Davis, R. L. Gene expression systems in Drosophila: a synthesis of time and space. Trends Genet 20, 384–391 (2004).

28. Talay, M. et al. Transsynaptic Mapping of Second-Order Taste Neurons in Flies by trans-Tango. Neuron 96, 783–795.e4 (2017).

29. Glicksman, M. A. & Brower, D. L. Expression of the Sex combs reduced protein in Drosophila larvae. Dev Biol 127, 113–118 (1988).

30. Liu, Z. et al. Par complex cluster formation mediated by phase separation. Nat Commun 11, 2266 (2020).

31. Risse, B. et al. FIM, a novel FTIR-based imaging method for high throughput locomotion analysis. PLoS One 8, e53963 (2013).

